# Microbiome variability of mosquito lines is consistent over time and across environments

**DOI:** 10.1101/2023.04.17.537119

**Authors:** Anastasia Accoti, Shannon Quek, Julia Vulcan, Cintia Cansado-Utrilla, Enyia R Anderson, Jessica Alsing, Hema P. Narra, Kamil Khanipov, Grant L. Hughes, Laura B. Dickson

## Abstract

The composition of the microbiome is shaped by both environment and host genetic background in most organisms, but in the mosquito *Aedes aegypti* the role of host genetics in shaping the microbiome is poorly understood. Previously, we had shown that four lines of *Ae. aegypti* harbored different microbiomes when reared in the same insectary under identical conditions. To determine whether these lines differed from each other across time and in different environments, we characterized the microbiome of the same four lines of *Ae. aegypti* reared in the original insectary and at another institution. While it was clear that the environment influenced the microbiomes of these lines, we did still observe distinct differences in the microbiome between lines within each insectary. Clear differences were observed in alpha diversity, beta diversity, and abundance of specific bacterial taxa. To determine if the line specific differences in the microbiome were maintained across environments, pair-wise differential abundances of taxa was compared between insectaries. Lines were most similar to other lines from the same insectary than to the same line reared in a different insectary. Additionally, relatively few differentially abundant taxa identified between pairs of lines were shared across insectaries, indicating that line specific properties of the microbiome are not conserved across environments, or that there were distinct microbiota within each insectary. Overall, these results demonstrate that mosquito line can shape the microbiome across microbially- diverse environments and host by microbe interactions affecting microbiome composition and abundance is dependent on environmentally available bacteria.

**Author Summary:** The mosquito microbiome plays a critical role in shaping interactions with human pathogens. The factors that contribute to shaping the composition of the mosquito microbiome are of high importance due to its role in pathogen interactions and the successful development of control strategies. In other organisms, both host genetics and environment shape the microbiome composition, but the role of host genetics in shaping the mosquito microbiome is less clear. Previously, we have shown that different lines of *Aedes aegypti* harbor different microbiomes when reared in the same environment. We were curious to see if these differences could still be detected after further generations in the same insectary and across environments in a different insectary. We found that found that the microbiome differed between these lines in each insectary indicating an element of both host genetic background and environment play a role in establishing the microbiome. Our results indicate that different genetic backgrounds of *Ae. aegypti* will interact with their environment differently to shape their microbiome, which could potentially influence interactions with human pathogens and/or the effectiveness of control strategies. More broadly, our results are of interest for the ecology of host-microbe interactions.

## Introduction

The microbiome plays a crucial role in the health of various organisms [1, 2]. Alterations in the relative abundance and overall bacterial community structure can lead to dysbiosis within the organism resulting in disease or morbidity. Factors that shape the acquisition and maintenance of the microbiome vary between organisms and include contributions from both the environment and the host. The role of host genotype in shaping the composition of the microbiome has become apparent in mammalian systems where specific host genomic loci have been associated with specific bacterial taxa [3–5]. Additionally, the role of host genetic background in invertebrates such as *Drosophila* and other insects has also been shown to contribute to the composition of the microbiome [6–9]. While the contribution of host genetic background in shaping the microbiome is well studied in mammalian and *Drosophila* systems, few and contradictory data exist for the role of host genetic background in shaping the microbiome of the mosquito [10–12].

*Aedes aegypti* is the main vector of arthropod-borne viruses (arboviruses) worldwide, such as dengue, Zika, and chikungunya. Examples such as dengue virus (DENV) result in 100-400 million infections annually and remain a major threat to public health [13, 14]. The microbiome, specifically the presence of distinct isolates, plays a role in the ability of *Ae. aegypti* to be a successful vector of human pathogens [15, 16]. The microbiome of *Ae. aegypti* is largely shaped by the environment [17, 18], but the role of host genetic background remains poorly understood. Multiple studies have controlled for environmental variation and characterized the microbiome of diverse lines of *Ae. aegypti* reared in the same insectary (i.e. the same environment). These studies have found contradictory results. Two independent studies identified differences in the microbiome from the whole body of *Ae. aegypti* that were dependent on mosquito line [10, 11]. A third study did not observe any differences in either the bacterial community structure or the abundance of specific taxa in the midgut from a selection of genetically diverse lines of *Ae. aegypti*, which represented their worldwide genetic diversity [12]. These contradictory results suggest that perhaps the environment is important for detecting line specific differences in the microbiome, or alternatively, there are host and environmental interactions that determine the microbiome composition in *Ae. aegypti*.

*Aedes aegypti* occupies a variety of environments worldwide, allowing for the association of numerous bacterial taxa with it. Understanding the relative contributions of *Ae. aegypti* genetic background and the environment in shaping the microbiome composition is important for both teasing apart the role of the microbiome in mosquito vectorial capacity, [15] as well as development of paratransgenic control tools [19–21]. To understand whether line specific differences in the microbiome is dependent on the environment, we sequenced the 16s ribosomal RNA gene from four lines of *Ae. aegypti* that previously showed line specific differences in the microbiome [10] and the same four lines after being transferred to a new insectary at another institution. We analyzed the structure of the bacterial community between and within each insectary, as well as the differential abundance of specific taxa. Finally, we identify conserved genera that differ in pairwise comparison between lines at each insectary. Our findings that lines harbor differences in their bacterial composition despite being reared under identical conditions has important implications when comparing phenotypic effects which may be sensitive to the microbiome.

## Materials and Methods

### Ae. aegypti Mosquitoes

Colonies of *Ae. aegypti* used in this study originated from Galveston, Iquitos, Juchitan, and Thailand. At UTMB, the generation of the colonies is not known but the colonies have been in the insectary since 2010 and reared continuously. The colonies were transferred to LSTM in 2018. At each institute the mosquito lines were housed under standard insectary conditions consisting of 28°C and 70% +/− 10% relative humidity with 12h:12h light dark cycle. Eggs were hatched in deionized water and larvae were fed fish food. Adults were held in Bugdorm cages with constant access to 10% sucrose until being harvested.

### DNA Extractions

Three-five days post emergence, *Ae. aegypti* were cold anesthetized and females taken for surface sterilization. Individual mosquitoes were surface sterilized in ethanol 70% for 5 minutes followed by 3 washes in sterile PBS. DNA was extracted from 20 female mosquitoes from Galveston, Iquitos, Juchitan and Thailand using the QIAamp DNA Mini Kit (QIAGEN) following the manufacturer protocol with the following modifications: initial volume of 180ul of buffer ATL used for mosquito homogenization and final volume of 100ul nuclease-free water for DNA elution. No RNase A treatment was applied. For the UTMB samples, DNA from mosquitoes from Iquitos and Juchitan was extracted on 25 February 2021 and DNA from mosquitoes from Galveston and Thailand was extracted on 26 February 2021. No-mosquito controls were used for each extraction batch and sequenced. The negative control clustered differently from the samples (Supplemental Figure 1), with the exception of Juchitan.

### Library Preparation and Sequencing

Sequencing libraries for each isolate were generated using universal 16S rRNA V3-V4 region primers [22] in accordance with Illumina 16S rRNA metagenomic sequencing library protocols. DNA concentrations of each library were determined by Qubit and equal amounts of DNA from each barcoded library were pooled prior to sequencing. The samples were barcoded for multiplexing using Nextera XT Index Kit v2. The pooled libraries were diluted to 4 pM and run on the Illumina Miseq using a MiSeq Reagent Kit v2 (500-cycles).

### Data Analysis

To identify known bacteria, sequences were analyzed using the CLC Genomics Workbench 21.0.5 Microbial Genomics Module (CLC MGM). Reads containing nucleotides below the quality threshold of 0.05 (using the modified Richard Mott algorithm) and those with two or more unknown nucleotides or sequencing adapters were trimmed out. Reference-based Operational Taxonomic Unit (OTU) picking was performed using the SILVA SSU v132 97% database [23]. Sequences present in more than one copy but not clustered to the database were placed into de novo OTUs (97% similarity) and aligned against the reference database with an 80% similarity threshold to assign the “closest” taxonomical name where possible. Chimeras were removed from the dataset if the absolute crossover cost was three using a k-mer size of six. OTUs with a combined abundance of less than two were removed from the analysis. Low abundance OTUs were removed from the analysis if their combined abundance was below 10 or 0.1% of reads. Differential abundance analysis was performed using CLC MGM at the genus level to compare the differences between the groups using trimmed mean of M-values. Each OTU was modeled as a separate generalized linear model, where it is assumed that abundances follow a negative binomial distribution. The Wald test was used to determine significance between groups. Tables of differentially abundant taxa are on the complete unfiltered data set (Supplemental Tables 1 and 2). To perform the analyses of the number of genera differentially abundant shared between various pairwise comparisons, the ID of the genera from the filtered OTU table were pulled out of the full data file using the “join” command in R. Only the ID of genera from the filtered OTU table were used.

Abundance profiling was performed using MicrobiomeAnalyst [24, 25]. The analysis parameters were set so that OTUs had to have a count of at least 10 in 20% of the samples and above 10% inter-quantile range. Analysis was performed using actual and total sum scale abundances. Alpha diversity was measured using the observed features to identify the community richness using Chao1. Statistical significance was calculated using T-test/ANOVA. Beta diversity was calculated using the Bray-Curtis dissimilarity measure (genus level). Permutational Multivariate Analysis of Variance (PERMANOVA) analysis was used to measure effect size and significance on beta diversity for grouping variables [26]. Relative abundance analysis was done in MicrobiomeAnalyst at the level of genera.

### Upset plot

To identify the distribution of bacteria taxa between the different mosquito lines and their rearing insectaries, count data from female mosquitoes were extracted and aggregated to the genus level for each sample. This yielded a maximum of 44 distinct bacterial taxa (including ‘uncultured’ and ‘ambiguous’ taxa). This table of count data was then processed using the R package UpsetR [27], stratifying the data based on mosquito strain and the rearing insectary.

### Correlation plot

Previous studies have indicated a potential correlation in presence/absence between specific genera of bacteria-namely *Cedecea*, *Enterobacter*, *Klebsiella*, and *Serratia*. To test for this, counts from the four genera were extracted from the aggregated table described earlier. Using the sum of these counts as the total, percentage relative abundance for each taxa was then counted per individual sample. These results were then passed to the cor() function in R’s stats package [28] to obtain a correlation matrix of size four-by-four. This correlation matrix was then visualised using the R package corrplot [29], with the two options method = “color”, col = COL2(‘PiYG’).

## Results

### Between Insectary Differences

We previously reported that the microbiomes of four *Ae. aegypti* lines reared in the same insectary were different [10]. To confirm whether these differences were conserved across subsequent generations and environments, we characterized the microbiome of these lines in the insectary that they have been continuously reared in (UTMB) and an insectary at a different institution (LSTM) after being transferred and reared for multiple generations. To compare the microbiome diversity and composition between lines we undertook amplicon sequencing on the V4-V5 region of the bacterial 16s rRNA gene from 20 individuals from each line. Out of the 160 individuals sequenced, a total of 79 OTUs were found which represented 44 bacterial genera. Rarefaction curves (Supplemental Figure 1) show that sufficient sequencing depth was achieved.

To determine if the microbiome of each *Ae. aegypti* line differed in diversity of bacterial species present, the within line diversity was determined by calculating the Chao Diversity index (Figure 1A). The richness of the microbiome was greater in mosquitoes reared in the LSTM insectary compared to those reared in the UTMB insectary (p-value < 0.001). To determine if the community structure of any one line was more closely related to the same line reared at a different insectary, or different lines reared within the same insectary, principal component analysis (PCA) was performed on a Bray-Curtis dissimilarity matrix. Overall, individuals from each line shared a more similar bacterial community structure with other lines from the same insectary (Figure 1B), indicating that the environment and non line-specific factors plays a more influential role in shaping the community structure of the microbiome.

**Figure 1:**
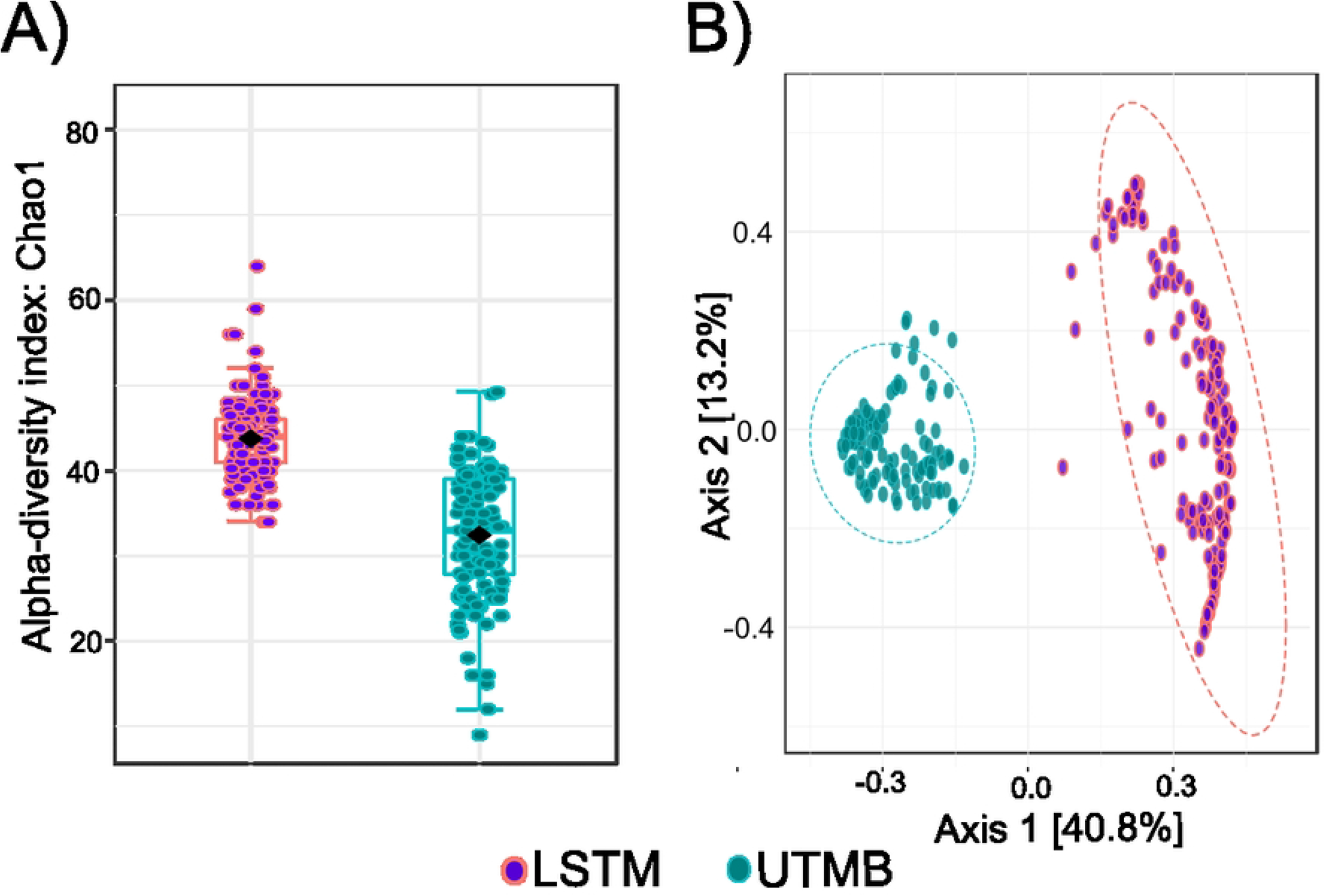
Diversity of the microbiome in individuals reared at the LSTM or UTMB insectaries. Structure of bacterial communities was determined by deep sequencing the V3-V4 region of the bacterial 16*S* gene in adults from 4 different lines of *Ae aegypti* (Galveston, Thailand, Iquitos, Juchitan) reared in two different insectaries at UTMB and LSTM. The Bacterial community structure is represented (A) by the species richness index Chao1 and (B) by principal component analysis of Bray-Curtis dissimilarity index. Mosquito lines reared at LSTM are shown in purple, mosquitoes reared to UTMB are shown in turquoise.

### Within Insectary Differences

To determine whether the lines differed from each other within the same insectary, alpha and beta diversity was compared between the lines from each insectary. Differences in species richness were observed between the lines at both the LSTM insectary (Figure 2A) (p-value < 0.001) and the UTMB insectary (Figure 2C) (p-value < 0.001). To gauge if the lines differed in the bacterial community structure, PCA was performed on a Bray-Curtis dissimilarity matrix. The PCA plots show that the bacterial communities differ between the lines at both the LSTM insectary (Figure 2B) (p-value: 0.001) and the UTMB insectary (Figure 2D) (p-value: 0.001). Strikingly, these results indicate that differences in the bacterial community structure of these four lines are maintained across generations and insectaries.

**Figure 2:**
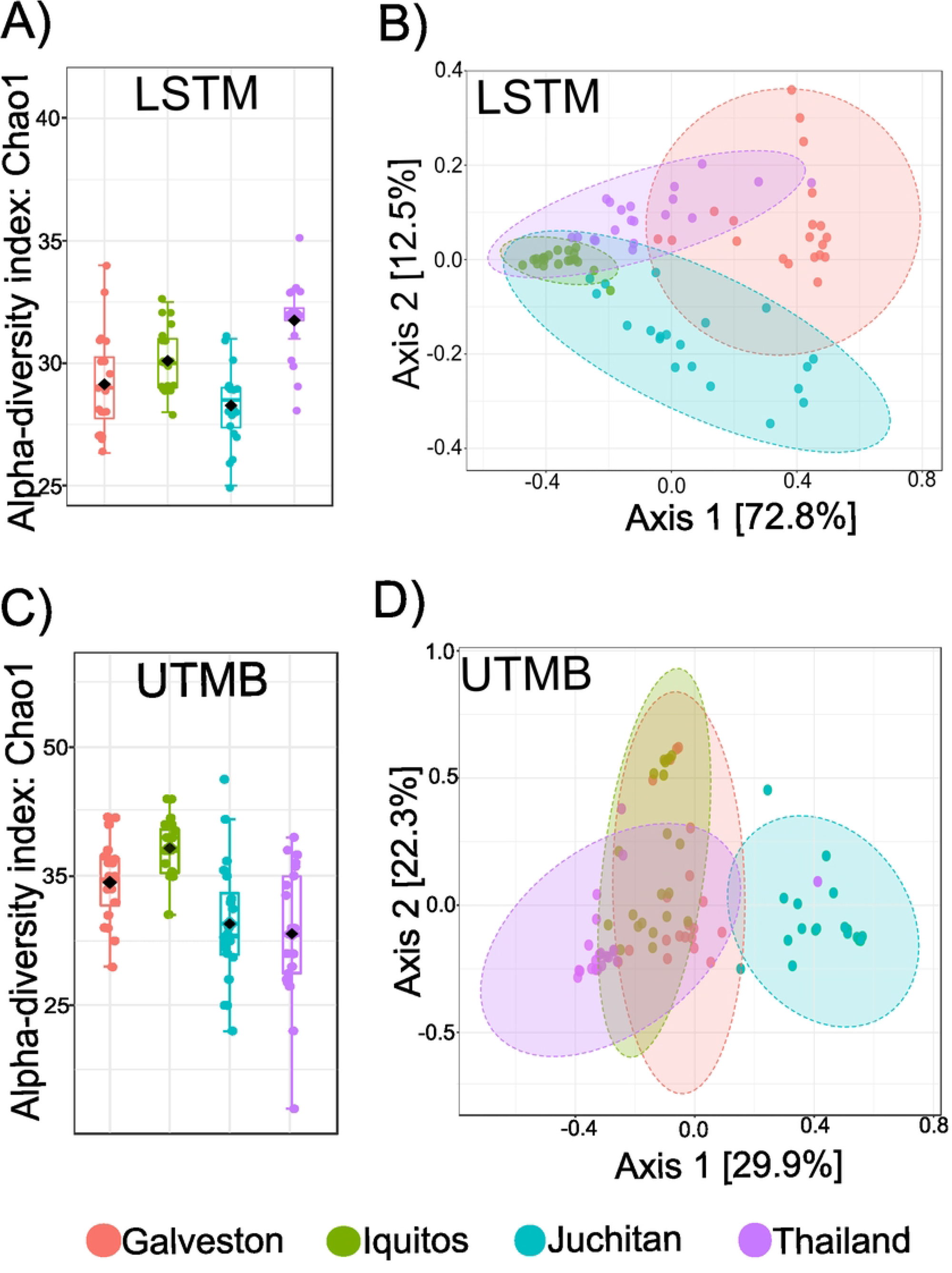
Alpha and beta diversity metrics for the *Ae. aegypti* lines reared at the LSTM and UTMB insectaries. (A, C): The species richness (Chao1 index) was calculated from 20 individuals from each line (Galveston, Thailand, Iquitos, Juchitan) at each insectary. The level of species richness differed between individuals from the LSTM (p-value < 0.001) and UTMB (p-value < 0.001). (B,D):The dissimilarities between the 4 different lines of *Ae. aegypti* was analyzed by principal component analysis of Bray-Curtis dissimilarity index. The bacterial community structure of the lines differed in individuals from LSTM (p-value = 0.001) and UTMB (p-value = 0.001).

To measure the extent of overlap of bacteria identified between the two insectaries, the presence of each genera was compared between each line from each insectary (Figure 3). Out of the 44 identified genera, 23 were present in both insectaries. Six genera are specific to the UTMB insectary, and three genera were specific to the LSTM insectary (Supplemental Table 3). We examined the relative abundance of the 20 most abundant genera for each line to determine if the bacterial communities were dominated by the same bacterial genera in each line at both insectaries. Different bacteria genera dominated mosquitoes reared in the LSTM insectary compared to the UTMB insectary (Figure 4). However, within both insectaries, the Juchitan lines showed the greatest difference in the relative abundance of different taxa compared to the other lines. All four lines reared in the LSTM insectary were dominated by an ambiguous taxa, *Perfucidibaca*, and *Chryseobacterium*. The four lines reared in the UTMB insectary had a high proportion of *Acinetobacter*, *Psuedomonas,* and *Asaia*.

**Figure 3:**
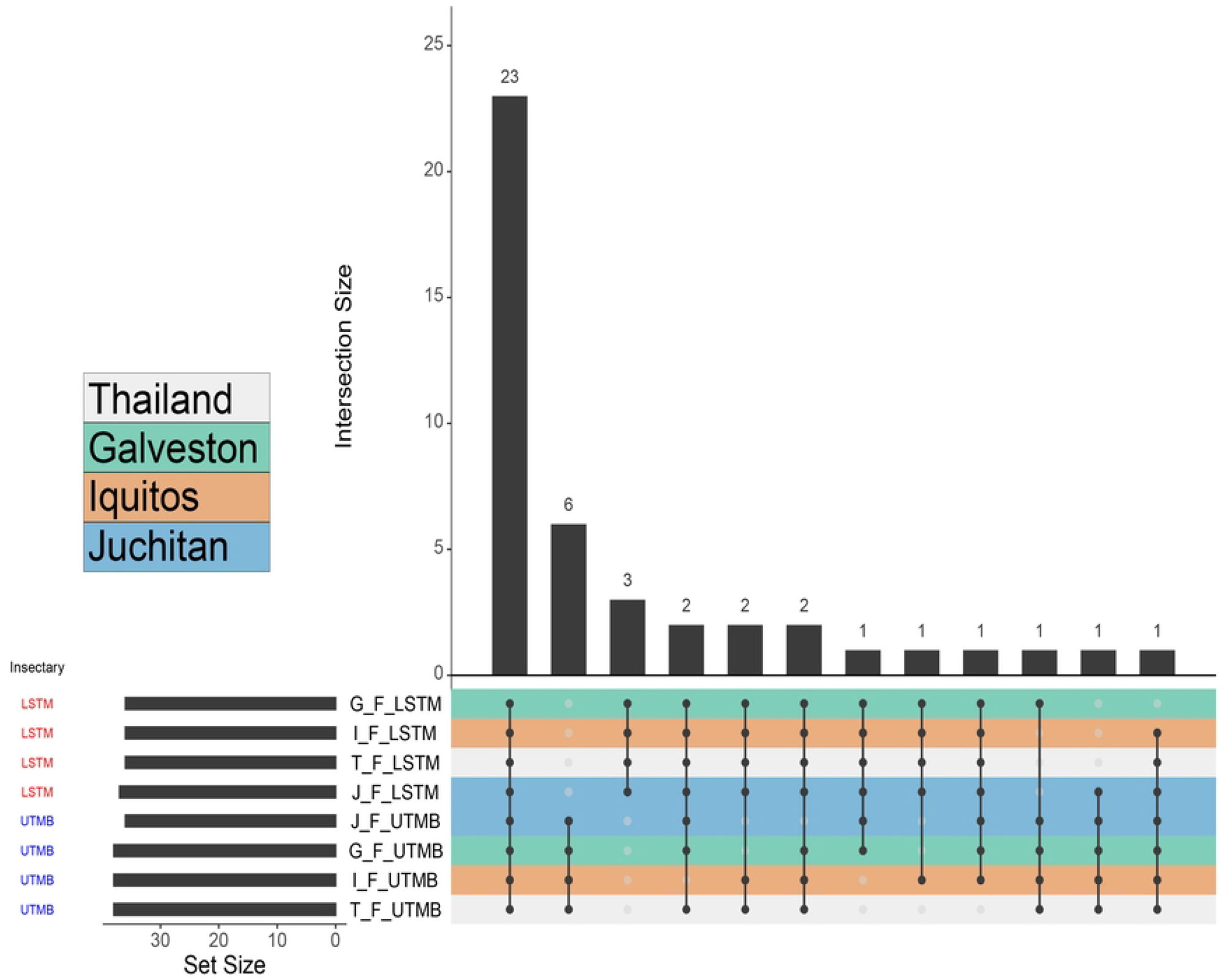
Upset plot showing the number of genera that are shared between lines in the LSTM and UTMB insectaries. Each mosquito strain in the different insectary (one per row at the bottom half of the image) is treated as a ‘set’ with an identified number of bacterial taxa (‘Set Size’). The various permutations of intersections are denoted by the ball-and-stick diagram at the bottom of the image, and size of these intersections denoted by the bar graph at the top of the image (‘Intersection Size’). Rows are colored based on mosquito strain (middle-left), and further divided into the rearing Insectary (bottom left). 23 taxa are shared across all the different strains between the different insectaries, constituting a potential ‘core’ set of bacteria. This is then followed by six (Persicitalea, Janthinobacterium, Rahnella, Uncultured, Luteolibacter, Verrucomicrobium) and three taxa (Sphingopyxis, Burkholderia-Caballeronia-Paraburkholderia, Methyloversatilis) that appeared unique to the rearing insectary. All other permutations of intersects contain two or fewer bacterial taxa.

**Figure 4:**
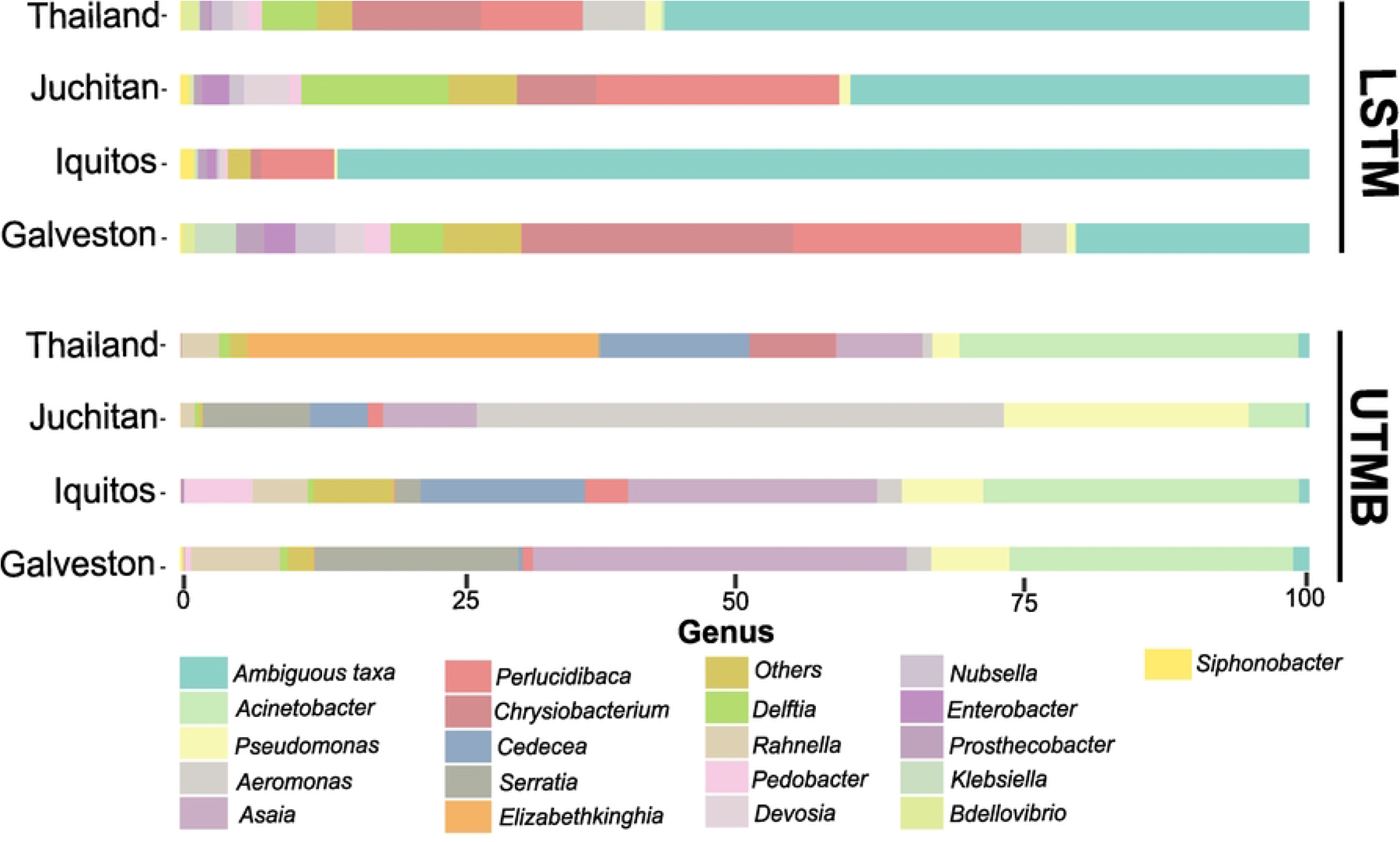
Relative abundance of bacteria in each line. The dominant bacterial genera are different between insectaries and between lines. The relative abundance of the 20 most abundant genera in shown for 20 individuals from each line at LSTM and UTMB insectaries. Bacterial genera were assigned to OTUs clustered with a 97% sequence identity cutoff and taxonomically classified with the SILVA database.

We previously reported a correlation between Enterobacteriaceae bacteria and *Serratia* in *Ae. aegypti* [10]. Specifically, if the midgut of *Ae. aegypti* was colonized with Enterobacteriaceae, *Serratia* was excluded from colonizing. The robustness of this phenotype after multiple generations and in different environments was measured. We found that this negative correlation between the presence of Enterobacteriaceae and the presence of *Serratia* was still observable in these lines (Supplemental Figure 2).

### Environment versus Line Specific Differences

To understand if line specific differences are conserved across environments, pair-wise differences in the abundance of each bacterial genera were calculated between lines within and across insectaries (Supplemental Tables 1 and 2). Out of the 44 genera identified, a range of 16-38 genera were found in different abundances between lines within each insectary resulting in 36-86% similarity between the lines (Figure 5). To assess whether line specific factors or the environment play a larger role in shaping the microbiome, the percentage of genera differentially abundant was compared for each pairwise comparison within and between insectaries. Lines were more similar to lines from the same insectary than to the matching line in the other insectary (Figure 5), indicating the environment is more important than line specific factors in shaping the microbiome.

**Figure 5:**
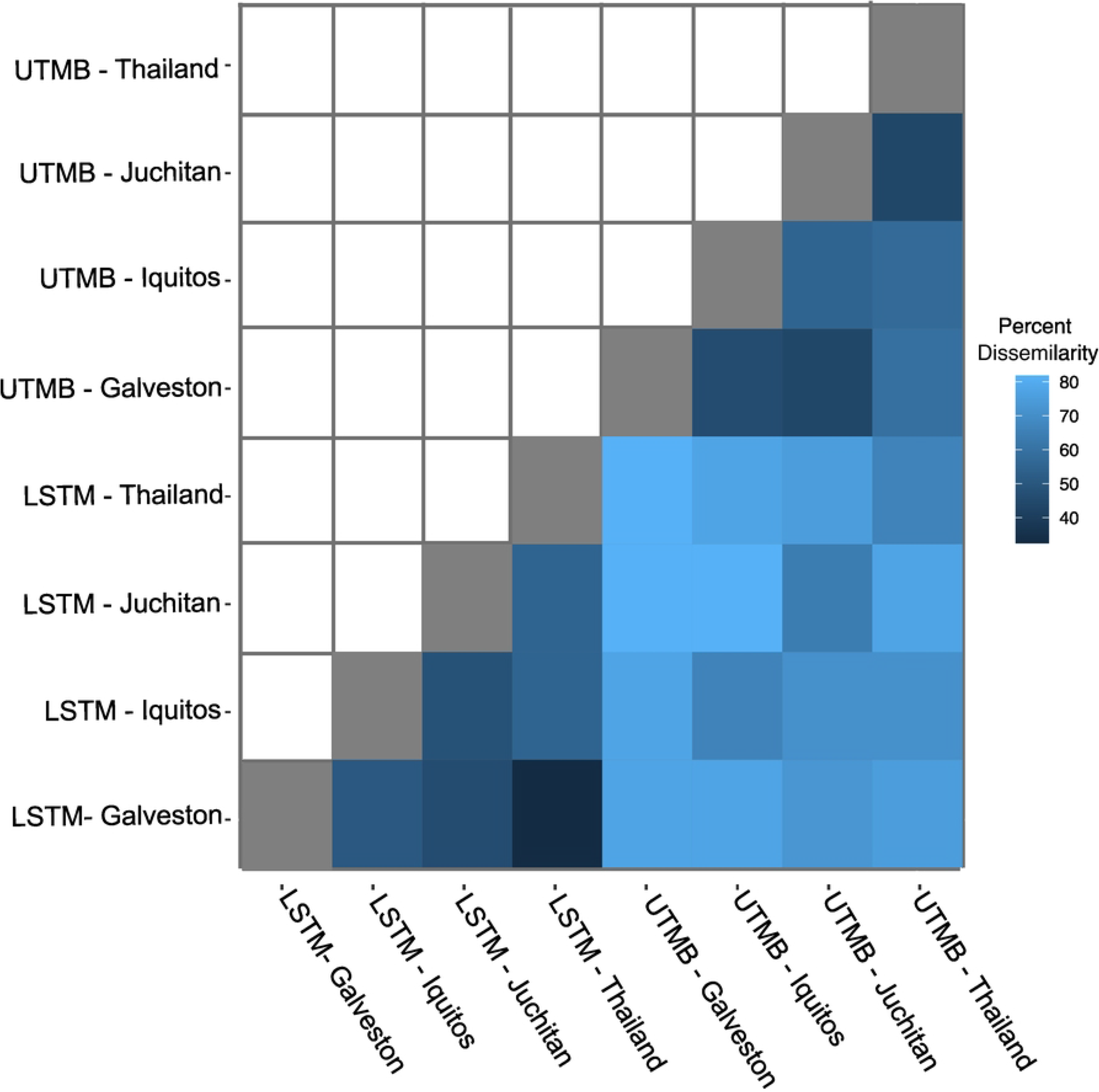
Pairwise comparisons of mosquito lines reared in each insectary. The microbiomes of mosquito lines reared in the same insectary are more similar compared to those reared in a different insectary. Results from a pairwise differential abundance analysis are shown for each pair of lines as the percent of genera that are significantly different between the pairs after correcting for multiple comparisons. A light blue color indicates a higher degree of dissimilarity between the lines.

To assess if lines differ from each other in the same way across environments, pairwise comparisons of lines were performed within each insectary, and genera that showed statistically differential abundances were identified per comparison. If we identify conservation of differentially abundant genera between the same two lines reared in different environments, this would suggest an interaction between specific bacterial genera and the hosts’ background across environments. Overall, a range of 11-19 genera were identified as having conserved differences in their abundance between the same lines in each insectary (Figure 6). The proportion of differentially abundant genera between lines that were shared between institutes is dependent on the specific pairwise comparison (Figure 6, Supplemental table 4). In pairwise comparisons between the Galveston vs Iquitos and Galveston vs Juchitan lines, roughly half of the differentially abundant bacteria are conserved across insectaries. In the other four pairwise comparisons, more than half of the differentially abundant bacteria are conserved across insectaries, indicating that the structuring of the microbiome between lines from the same environment is partially conserved across environments.

**Figure 6:**
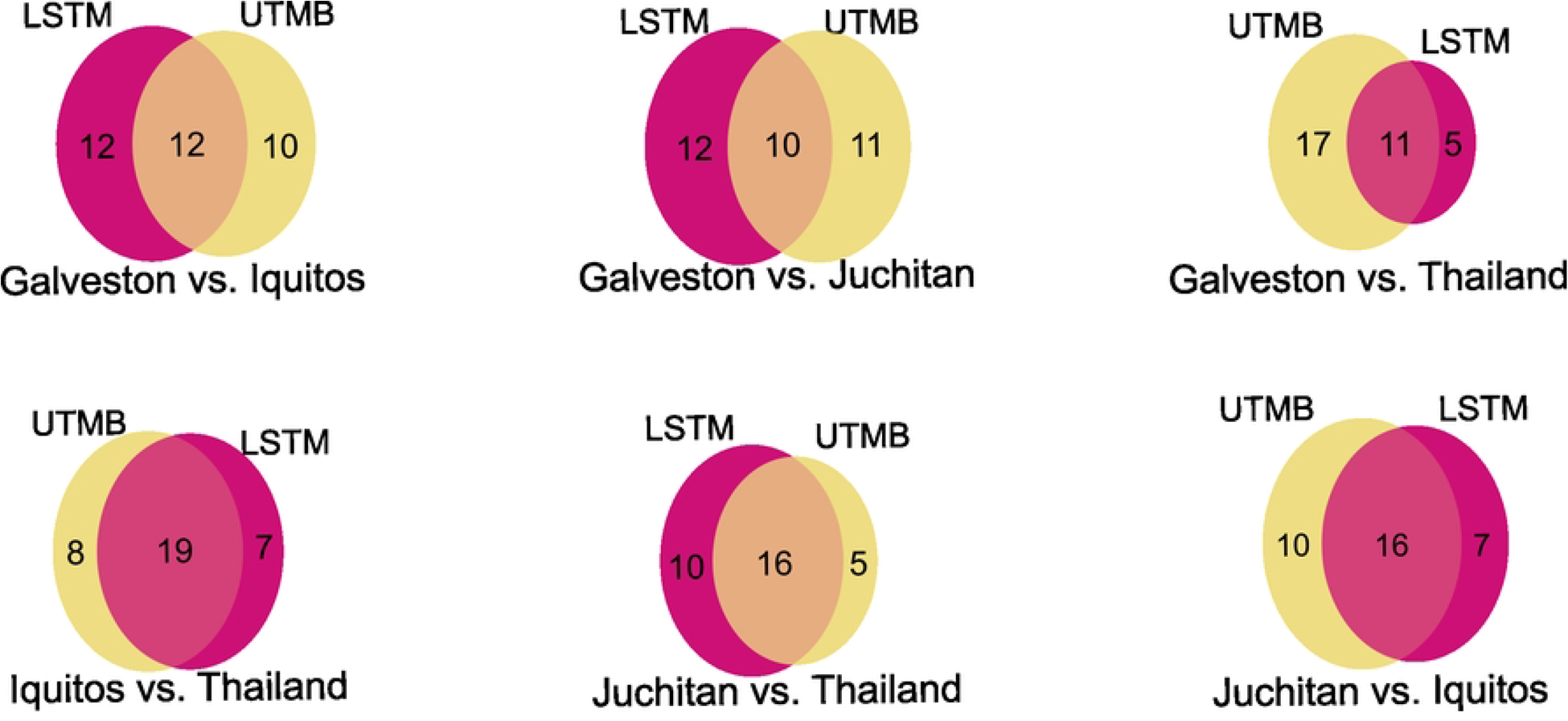
Few genera have conserved differences in abundance between lines at both insectaries. Venn diagrams show the overlap in specific genera that are differentially abundant between each pair of lines (Galveston, Iquitos, Thailand, or Juchitan) at each insectary (LSTM or UTMB). The ID of genera differentially abundant between each pairwise comparison was compared between insectaries. The shared genera represent a genus that is differentially abundant between the lines in both environments. The ID of shared genera can be found in supplemental table 4.

## Discussion

We explored the contribution of mosquito line and environment in shaping the microbiome composition of *Ae. aegypti*. Having previously observed differences in the microbiome between four lines of *Ae. aegypti* reared under identical conditions in the same insectary [10], we sought to determine if these lines continue to have distinct microbiomes since our previous work, and if this was consistent after rearing in different insectary environments. In accordance with our previous study [10], we observed differences in alpha and beta diversity between the four lines and interestingly, the microbiome of these lines were still distinct after rearing in separate insectaries. Notably, we found that although the microbiomes of the lines differed from each other within an insectary, any one line was more similar to other lines reared within the same insectary as compared to the same line at the other insectary. Finally, we observed a percentage of differentially abundant genera between lines were shared between insectaries. Together these data demonstrate that while the environment influences the microbiome available to the organisms, host factors still play a role in shaping the microbiome.

The role of host genotype in determining the microbiome composition has been established in other systems [3–9]. Although the influence of mosquito genotype in shaping the microbiome in the same environment is not well understood, multiple mosquito genes have been identified that influence the composition of the microbiome and gut homeostasis [11, 30–33]. Genes involved in bloodmeal digestion and immune factors can regulate the abundance of the microbiome as a whole [33], or the abundance of specific taxa [30]. Furthermore, the mosquito microbiome can stimulate expression of specific genes to shape the immune status of the mosquito allowing for efficient colonization of specific microbes [32]. In addition to immune genes regulating microbiome composition, metabolic signaling and nutrition status of the mosquito can influence the microbiome [11, 15]. Immunity and nutritional processes are under genetic control and different genotypes of *Ae. aegypti* could result in differences in levels of micro- and macronutrients, which in turn could influence the ability of specific bacteria to colonize that mosquito line. Additionally, genetic variation in immune genes could influence the success of different bacteria in colonizing the mosquito.

If mosquito line played a more profound role than the environment in determining which genera colonize the mosquito, we would expect the same line at the two insectaries to share more similarities to each other than the other lines within the same insectary, assuming similar taxa were available to the mosquito across insectaries. Additionally, this would be apparent if pairwise differences in taxa abundance between any two lines within one insectary were conserved in another insectary. We observed the opposite of this where lines within one insectary had more similar abundances of specific genera to each other, compared to the same line reared at a different insectary. We also observed that pairwise differences in the abundance of different genera between lines within one insectary were not shared in another insectary. This lack of shared differences in the abundance of specific genera within a line across institutes suggest that the environment, or the microbes the mosquito has access to, plays a larger role than host specific factors in determining the microbiome. Given that only 25 of the 44 genera identified were shared across insectaries, it is not surprising that line specific abundances of specific taxa are lost across insectaries.

This study and previous studies observed differences in the microbiome between the lines when reared under identical conditions in the same insectary [10]. This is contrast to our previous work in which midguts from diverse genotypes of *Ae. aegypti* harbored the same microbiome when reared under identical conditions [12]. The discrepancy between these studies could be a result of what tissues were used for microbiome characterization. Kozlova et al. [10] and this study sequenced the microbiome from the whole mosquito, while Dickson et al. [12] sequenced the midgut microbiome. Additionally, it is possible that the environmental bacteria present in the insectary in Dickson et al. [12] could colonize the midguts of all the lines effectively, while this was not the case in the UTMB and LSTM insectaries. Microbe-microbe interactions are important for determining the microbiome composition [17] and perhaps some of the environmental bacteria present in both the UTMB and LSTM insectaries are incapable of colonizing *Ae. aegypti*.

Given that the lines of *Ae. aegypti* were not sequenced in this study we cannot conclude these lines represent different genotypes of *Ae. aegypti*. The colonies used in the study represent a similar genetic background in relation to global populations of *Ae. aegypti* [34, 35]. Without sequencing the different lines to confirm genetic differences, we cannot unequivocally determine if the differences we see in the microbiome composition are correlated with mosquito genotype. Although we expect them to be similar genetically, it is likely that there is genetic divergence between the lines and it is reasonable to conclude that the differences between the microbiome between the lines is at least partially genetically controlled.

The results from this study provide a unique opportunity to tease apart host control of the microbiome in *Ae. aegypti*. By exploiting our new cryopreservation method [36] and swapping the microbiome using our recently developed transplantation approach [37] from both lines and insectaries, the role of the host in controlling the microbiome could be further investigated and provide insight into host genetic mechanisms that underly microbiome composition.

## Acknowledgements.

LBD was supported by UTMB start-up funds. GLH was supported by the BBSRC (BB/T001240/1, BB/V011278/1, BB/X018024/1, and BB/W018446/1), the UKRI (20197), a Royal Society Wolfson Fellowship (RSWF\R1\180013), the NIHR (NIHR2000907) and the Bill and Melinda Gates Foundation (INV-048598).

## Supplemental figures

**Supplemental Figure 1:** Rarefaction curves showing the sequencing depth of each library. The number of species is shown on the Y axis, and the number of sequencing reads is shown on the X axis.

**Supplemental Figure 2:** Heatmap showing the correlation between bacterial taxa co-occurrence between the different mosquito strains and rearing insectaries. Saturated greens indicate strong co-occurrence between the bacteria taxa, while saturated purple indicates strong exclusion, and whites indicate no positive or negative correlation. We show that *Klebsiella* co-occurs with *Enterobacter* taxa, whilst *Serratia* and *Cedecea* taxa appear to exclude other bacterial taxa.

**Supplemental Figure 3:** Beta diversity metrics for the *Ae. aegypti* lines reared at the UTMB insectary. The dissimilarities between the 4 different lines of *Ae. aegypti* plus the negative control was analyzed by principal component analysis of Bray-Curtis dissimilarity index.

